# Experimental SARS-CoV-2 infection using horseshoe bats

**DOI:** 10.64898/2026.03.19.712934

**Authors:** Kaoru Usui, Masumi Tsuda, Keiya Uriu, Shigeru Fujita, Yukie Kashima, Yutaka Suzuki, Lei Wang, Shinya Tanaka, Jumpei Ito, Kei Sato

## Abstract

Horseshoe bats are known as the natural reservoir of sarbecoviruses. To understand how horseshoe bats coexist with sarbecoviruses in nature, experimental infection can provide direct evidence. However, *in vivo* infection studies using horseshoe bats have been lacking because of the difficulty of maintaining insectivorous bats in a laboratory setting. Here, we established a stable husbandry system for greater horseshoe bats (*Rhinolophus ferrumequinum*) and performed experimental infection with SARS-CoV-2. In contrast to Syrian hamsters which showed substantial viral replication, infected horseshoe bats exhibited low-level but persistent viral replication in the lung without overt disease. Histological analyses revealed that inflammatory lesions in the bat lungs were spatially restricted and temporally delayed compared with those in hamsters. Transcriptomic analyses further showed preferential activation of tissue repair pathways but limited inflammatory responses following infection. Notably, bats expressed several interferon-stimulated genes prior to infection. Our results suggest that a host strategy combining constitutive antiviral state, limited inflammation and enhanced tissue repair may result in controlled viral replication without overt disease, likely enabling horseshoe bats to coexist with sarbecoviruses.

## Introduction

Sarbecoviruses (subgenus *Sarbecovirus*, genus *Betacoronavirus*) have caused two major outbreaks in humans over the past three decades: severe acute respiratory syndrome (SARS) caused by SARS-CoV and coronavirus disease 2019 (COVID-19) caused by SARS-CoV-2^1–3^. Multiple independent evidence from viral surveillance and phylogenetic analyses support horseshoe bats (family Rhinolophidae) as natural reservoir hosts of sarbecoviruses^4–10^. More broadly, bats harbor a remarkable diversity of zoonotic viruses, including filoviruses, henipaviruses, lyssaviruses, and coronaviruses^11,12^. These observations have fueled the hypothesis that bats have evolved host traits that promote disease tolerance, such as controlled inflammation and enhanced tolerance to tissue damage, allowing coexistence with a wide range of viruses with limited pathology^13,14^. Direct *in vivo* experimental infection in natural host–virus combinations is therefore essential to elucidate how bats coexist with viruses.

To date, *in vivo* infection studies in bats have been largely restricted to frugivorous species from the families Pteropodidae (for Hendra, Nipah, Marburg, and Ebola viruses) and Phyllostomidae (for influenza A virus), primarily owing to practical advantages in husbandry^15–24^. In contrast, horseshoe bats, the natural reservoir of sarbecoviruses, are insectivorous. Their strict dependence on live, flying insect prey makes long-term laboratory maintenance substantially more challenging. As a result, experimental systems for housing and infecting horseshoe bats under animal biosafety level 3 (ABSL-3) conditions have not been established. Surrogate approaches for sarbecoviruses, such as fruit bat models engineered to express human angiotensin-converting enzyme 2 (ACE2) in the lung, have reported persistent infection dynamics without visible clinical symptoms^25–28^. However, because these models do not reflect the natural host–virus relationship, both the antiviral and immunopathological responses of horseshoe bats to sarbecovirus infection, as well as the dynamics of sarbecovirus infection in their natural reservoir hosts, remain unaddressed.

Here, we first establish a stable experimental husbandry system for greater horseshoe bats (*Rhinolophus ferrumequinum*). By using this system, we conduct *in vivo* SARS-CoV-2 infection experiments under ABSL-3 conditions and characterize viral replication dynamics, pathology, and host responses, providing direct insight into the dynamics of sarbecoviruses infection in their natural reservoir hosts.

## Results

### Establishment of horseshoe bat husbandry system under ABSL-3 condition

Previous experimental infection studies in bats have been biased toward frugivorous species belonging to the families Pteropodidae and Phyllostomidae because they can be readily maintained as laboratory colonies^15–24^ (**Fig. 1A**). In contrast, horseshoe bats are insectivorous and feed exclusively on flying insects such as moths and flies in the wild (**Fig. 1A**). Their strict dietary dependence and high metabolic demands long been major obstacles to long-term maintenance, particularly under higher containment conditions.

**Figure 1.**
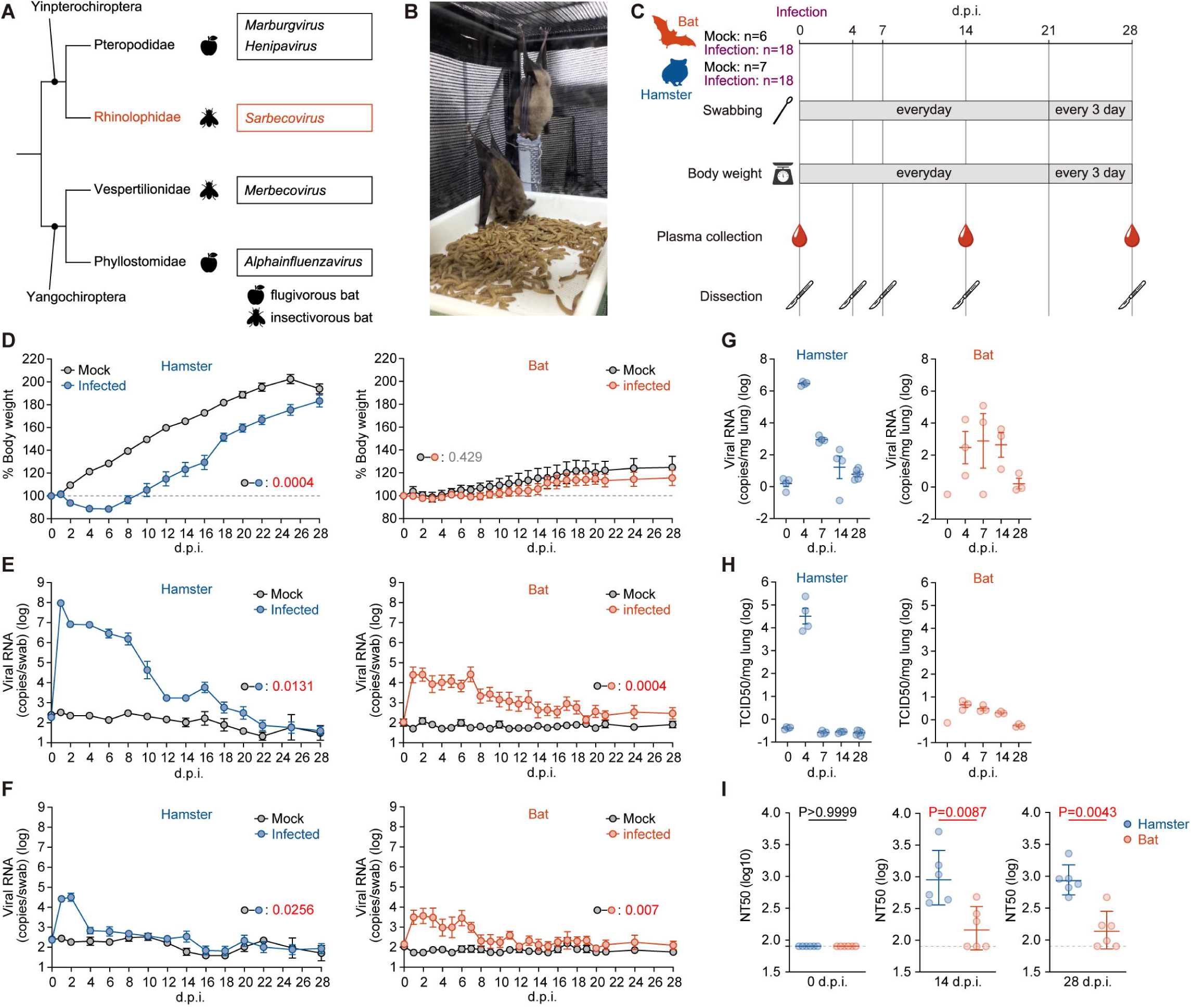
Infection dynamics of SARS-CoV-2 in greater horseshoe bats. Syrian hamsters (mock, n = 7; infected, n = 18) and greater horseshoe bats (mock, n = 12; infected, n = 18) were intranasally inoculated with SARS-CoV-2 Delta Q493K. **A**, Simplified phylogeny of bats (order Chiroptera), indicating representative virus genera hosted by each family and their dietary habits. The present study focuses on horseshoe bats (family Rhinolophidae), natural reservoir hosts of sarbecoviruses including SARS-CoV-2 (highlighted in red). **B**, Wild-captured greater horseshoe bats acclimated to self-feeding on mealworms in mesh cages within an ABSL-3 facility. **C**, Experimental design and sampling schedule. Oral and anal swabs and body weight were monitored daily until 21 d.p.i. and every 3 days thereafter until 28 d.p.i. Plasma samples were collected at 0, 14, and 28 d.p.i. Necropsies were performed at 0, 4, 7, 14, and 28 d.p.i. **D**, Percentage body weight change over experimental time. **E–F**, Viral RNA loads in oral swabs (**E**) and anal swabs (**F**) quantified by RT–qPCR. **G**, Viral RNA loads in lung tissues at 0, 4, 7, 14, and 28 d.p.i. **H**, Infectious virus titers in lung tissues measured by TCID50 assay. **I**, Neutralizing activity of plasmas against SARS-CoV-2 Delta Q493K spike-mounted pseudotyped virus at 0, 14, and 28 d.p.i., determined by pseudovirus neutralization assay. Data are shown as mean ± s.e.m. In **D–E**, statistical significance was cliculated by permutation test. In **I**, statistical analysis was performed by Wilcoxon signed-rank test.

In this study, greater horseshoe bats (*R. ferrumequinum)* were captured from caves in Japan and used to establish an experimental infection model. Based on previously reported husbandry methods developed for *R. cornutus*^29^, we adapted these approaches for the captured greater horseshoe bats and established a stable ABSL-3 husbandry system by training bats to self-feed on mealworms (**Fig. 1B** and **Supplementary Video 1**). Although individual variation was observed, most bats learned to self-feed within 1–2 weeks through a stepwise training protocol. After acclimating self-feeding, bat body weight started to increase steadily (**Supplementary Fig. 1**). Using this approach, horseshoe bats were stably maintained under ABSL-3 conditions for approximately six months until humane euthanasia according to the experimental design, demonstrating the feasibility of longer-term husbandry of wild-caught horseshoe bats by our husbandry system.

### Infection dynamics of SARS-CoV-2 in horseshoe bats

To establish an experimental infection model of horseshoe bats, we first selected SARS-CoV-2 rather than bat-derived coronaviruses, as its infection dynamics and pathogenicity have been extensively characterized in animal models^30–32^. Among SARS-CoV-2 variants, the Delta variant was selected because of its relatively higher pathogenicity compared with other variants^31^.

Consistent with previous report, pseudovirus-based entry assays showed that the Q493K substitution in the spike protein enables efficient utilization of angiotensin-converting enzyme 2 (ACE2) of *R. ferrumequinum* for viral entry^33^ (**Supplementary Fig. 2B**). Based on these findings, we artificially generated the replication-competent SARS-CoV-2 Delta variant bearing Q493K substitution in the spike protein (S:Q493K) by reverse genetics. Live virus infection assays showed that the Delta S:Q493K exhibited significantly higher infectivity in HOS-T cells expressing *R. ferrumequinum* ACE2 compared with the parental Delta strain (**Supplementary Fig. 2C**), validating its suitability for our *in vivo* infection experiment to *R. ferrumequinum*.

To investigate SARS-CoV-2 infection dynamics in horseshoe bats, these bats were intranasally inoculated with SARS-CoV-2 Delta S:Q493K at a dose of 10^5^ TCID50 under anesthesia and monitored for up to 28 days post infection (d.p.i.) (**Fig. 1C**). Prior to infection, we screened all bats using pan-sarbecovirus RT-PCR assay^34^ and serological testing against SARS-CoV-2 RBD reactivity, and confirmed that all bats included in this study were negative for active sarbecovirus infection and showed no evidence of prior sarbecovirus exposure (i.e., no detectable antibody against SARS-CoV-2 RBD). For the control of this experiment, we used Syrian hamsters (*Mesocricetus auratus*), which have been frequently used as an animal model for SARS-CoV-2 infection^35,30–32,36–38^, in the same experimental procedure we used for horseshoe bats. Consistent with previous report of experimental infection of Delta variant to hamster^36^, infected hamsters decreased body weight from 2 d.p.i., reaching a nadir at 4 d.p.i., and then recovered (**Fig. 1D**). Body weight changes in infected hamsters were significantly different from those in mock-infected controls (permutation test, P = 0.0004; **Fig. 1D**). In contrast, infected horseshoe bats did not exhibit statistically significant body weight loss compared with mock-infected controls (permutation test, P = 0.429; **Fig. 1D**). We then routinely quantified viral RNA collecting oral and anal swabs from infected animals. In oral swabs, viral RNA levels in infected hamsters peaked at 1 d.p.i. (**Fig. 1E**). On the other hand, horseshoe bats showed approximately 1000-fold lower levels of viral RNA compared to hamsters at 1 d.p.i. (**Fig. 1E)**. Viral RNA levels in horseshoe bats remained stagnant at a low level relative to hamsters until 18 d.p.i. In rectal swabs, while viral RNA was detected in hamsters only at 1 and 2 d.p.i., horseshoe bats showed persistent detection of viral RNA in rectal swabs up to 7 d.p.i. (**Fig. 1F**).

We next assessed the viral load dynamics in the lung at both the RNA and infectious virus levels. Viral RNA in lung tissues peaked in hamsters at 4 d.p.i. and subsequently declined between 7 and 28 d.p.i. (**Fig. 1G**). On the other hand, viral RNA in horseshoe bats was consistently detected at relatively low levels, averaging approximately 2–3 log_10_ copies per mg of lung tissue across 4, 7, and 14 d.p.i., without a clear peak (**Fig. 1G**). Quantification of infectious virus titer in lung tissues detected infectious virus only at 4 d.p.i. in hamsters (**Fig. 1H**). In bats, infectious viral titer was detectable in horseshoe bats across 4, 7, and 14 d.p.i. but was markedly low (**Fig. 1H**).

We then assessed the development of neutralizing antibody responses following infection. Our assay revealed that the plasma of infected horseshoe bats at 14 and 28 d.p.i. showed 50% neutralization titer (NT50) against SARS-CoV-2 Delta S:Q493K (**Fig. 1I**). However, these NT50 values were 10-fold lower than those of hamsters (**Fig. 1I)**.

### Lung pathology and immune responses to SARS-CoV-2 infection in horseshoe bats

To further investigate the differences of viral spreading in the lungs of the infected bats and hamsters, formalin-fixed right lungs of infected animals were analyzed at 4, 7, 14 and 28 d.p.i. We carefully identified the lobules and lobar bronchi sectioning each lobe along with the bronchial branches (**Fig. 2**).

**Figure 2.**
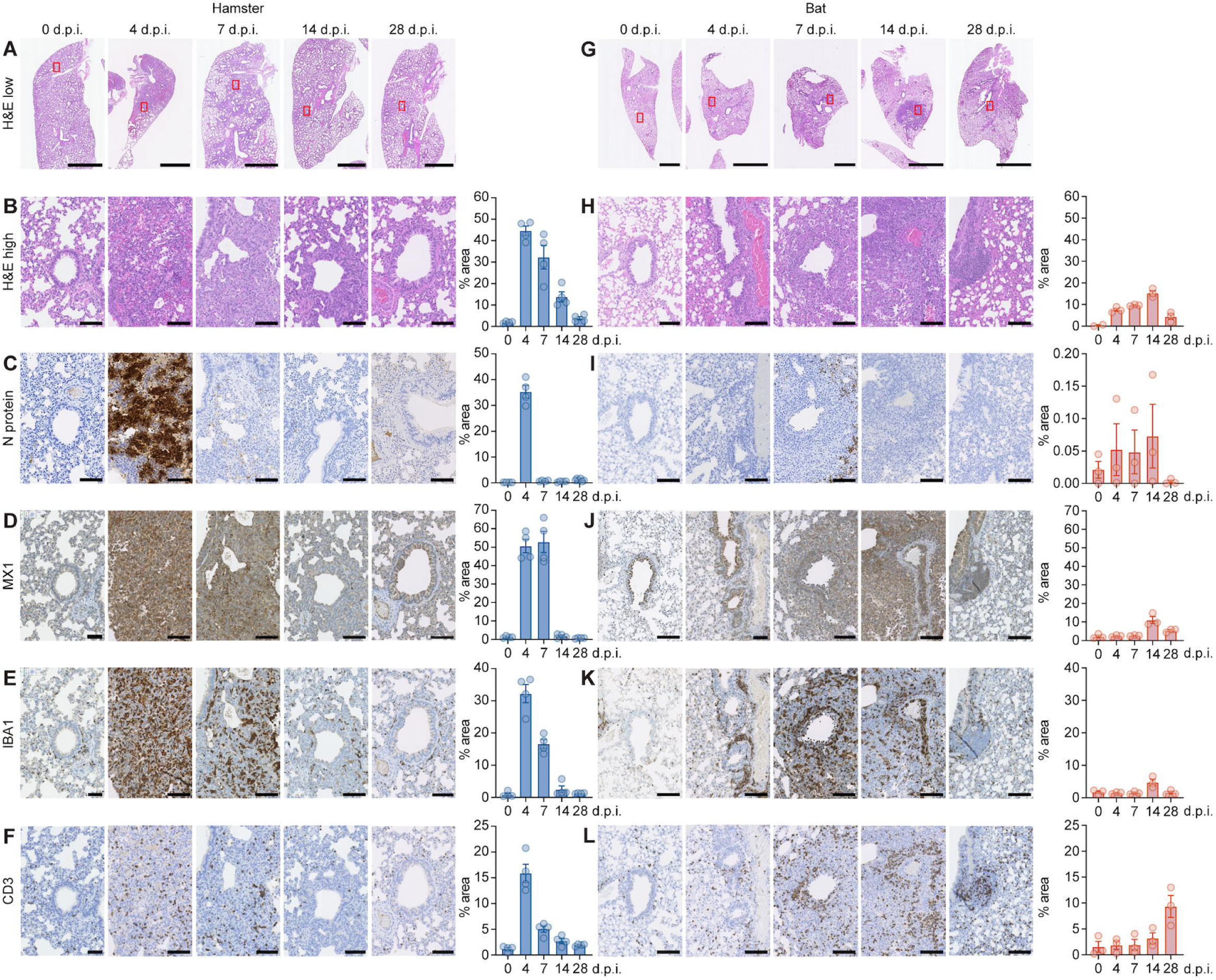
Histopathological features of SARS-CoV-2 in lung of greater horseshoe bats. Syrian hamsters (mock, n = 4; infected, n = 18) and greater horseshoe bats (mock, n = 3; infected, n = 12) were intranasally inoculated with SARS-CoV-2 Delta Q493K and subjected to histopathological analysis at the indicated time points. **A**, Representative H&E staining of lung sections from infected hamsters. Red boxes indicate regions shown at higher magnification in **B–F** for each timepoint. **B**, High-magnification H&E images and quantification of lesion area in hamster lungs. **C**, IHC for SARS-CoV-2 nucleocapsid (N) protein and quantification of N-positive area in hamster lungs. **D,** IHC for MX1 and quantification of MX1-positive area in hamster lungs. **E**, IHC for IBA1 and quantification of IBA1-positive area in hamster lungs. **F**, IHC for CD3 and quantification of CD3-positive area in hamster lungs. **G**, Representative H&E staining of lung sections from infected bats. Red boxes indicate regions shown at higher magnification in **H–L** for each timepoint. **H**, High-magnification H&E images and quantification of lesion area in bat lungs. **I**, IHC for SARS-CoV-2 N protein and quantification of N-positive area in bat lungs. **J**, IHC for MX1 and quantification of MX1-positive area in bat lungs. **K**, IHC for IBA1 and quantification of IBA1-positive area in bat lungs. **L**, IHC for CD3 and quantification of CD3-positive area in bat lungs. Data are presented as mean ± s.e.m.; each dot represents one individual animal. Scale bars, 2.5 mm (**A**, **G**) and 100 μm (**B–F**, **H–L**).

In hamsters, hematoxylin and eosin (H&E) staining demonstrated that inflammatory lesions paralleled viral kinetics, peaking at 4 d.p.i. and resolving thereafter (**Fig. 2A, 2B and Supplementary Fig. 4A**). At the 4 and 7 d.p.i., the lesion area was spread almost whole lung area **(Supplementary Fig. 4A)**. We then performed immunohistochemistry (IHC) as performed in our previous studies^36^. IHC of the SARS-CoV-2 nucleocapsid (N) protein in hamster (**Fig. 2C**) closely mirrored viral RNA levels and infectious titers in the lung **(Fig. 1G and 1H**). N-positive cells peaked at 4 d.p.i. and declined sharply thereafter (**Fig. 2C and Supplementary Fig. 5A**). At peak infection, viral antigen was ubiquitously distributed throughout the lung parenchyma (**Fig. 2C and Supplementary Fig. 5A**). IHC of MX1, a protein upon viral infection and type I interferon (IFN-I) stimulation, showed that hamsters exhibited robust MX1 expression at 4 and 7 d.p.i. (**Fig. 2D and Supplementary Fig. 6A**). The MX1 expression observed in a broad area in the lung parenchyma and became nearly undetectable by 14 and 28 d.p.i. (**Fig. 2D and Supplementary Fig. 6A**). IHC of IBA1, a marker protein of macrophages, in hamsters showed the rapid accumulation at 4 d.p.i. and declined thereafter corresponding to dynamics of inflammation lesion (**Fig. 2E and Supplementary Fig. 7A**). By 28 d.p.i., few IBA1-positive cells remained in the lung (**Fig. 2E and Supplementary Fig. 7A**). CD3-positive T cells in hamsters also rapidly accumulated in the lung at 4 d.p.i., coinciding with peak inflammation, and subsequently declined after 7 d.p.i. (**Fig. 2F and Supplementary Fig. 8A**).

In bats, H&E staining showed that the peak of inflammation lesion was at 14 d.p.i. (**Fig. 2G, 2H and Supplementary Fig. 4B**), which was profoundly later than that of hamsters (**Fig. 2A and 2B**). In contrast to hamsters, which showed a sharp increase in lesion area, bats exhibited a more moderate increase. Additionally, in contrast to hamster, the lesion area in the bats was spatially restricted around bronchiole (**Fig. 2G and Supplementary Fig. 4B**). IHC of SARS-CoV-2 N-protein showed that the number of N-positive cells was markedly lower compared to hamsters (**Fig. 2I and Supplementary Fig. 5B**). Although the number of N-positive cells was extremely low, N-protein signals were persistently detected from 4 to 14 d.p.i. (**Fig. 2I and Supplementary Fig. 5B**), which reflects the low-level viral replication observed in bats (**Fig. 1G and 1H**). Bats showed the peak of MX1-positive area at 14 d.p.i, which coincides with the delayed inflammatory response compared to hamsters (**Fig. 2J and Supplementary Fig. 6B**). At 14 d.p.i., the MX1-positive area was overlapped with the inflammatory lesion area (**Fig. 2H, 2J and Supplementary Figs. 5B and 6B**). Notably, MX1 expression was observed in the bronchial epithelium at all time points including uninfected 0 d.p.i. specimens (**Fig. 2J and Supplementary Fig. 6B**). IBA1-positive macrophages were accumulated at bronchi at 14 d.p.i. (**Fig. 2K and Supplementary Fig. 7B**). Then, these IBA1-positive macrophages infiltrated into inflammatory lesion area during 7–14 d.p.i. (**Fig. 2H, 2K and Supplementary Figs. 5B, 7B**). The number of CD3-positive T cells was gradually increased reaching a maximum at 28 d.p.i (**Fig. 2L and Supplementary Fig. 8B**). During 7–14 d.p.i., T cells accumulated at the periphery of inflammatory lesions area (**Fig. 2L and Supplementary Fig. 8B**). Altogether, these histological analyses suggest that the dynamics of SARS-CoV-2 infection and subsequent immune and inflammatory responses are profoundly distinct between bats and hamsters.

### Transcriptomic dynamics in the lung of SARS-CoV-2-infected horseshoe bats

To investigate the gene expression dynamics of horseshoe bats after SARS-CoV-2 infection, RNA sequencing (RNA-seq) was performed using the lung tissues collected at 0, 4 (inflammation peak in infected hamsters; **Fig. 2C**), and 14 (inflammation peak in infected bats; **Fig. 2I**) d.p.i. Differentially expressed genes (DEGs) were identified by comparing infected timepoints (4 and 14 d.p.i.) with mock samples (0 d.p.i.) in hamsters and bats respectively. In hamsters, the number of DEGs was higher at 4 d.p.i. than at 14 d.p.i. (**Fig. 3A**). In contrast, horseshoe bats exhibited a greater number of DEGs at 14 d.p.i. than at 4 d.p.i. (**Fig. 3A**). These differences in peak of DEGs number between hamsters and bats are consistent with temporal dynamics of inflammation in the lung respectively (**Figs. 2C,I and 3A**).

**Figure 3.**
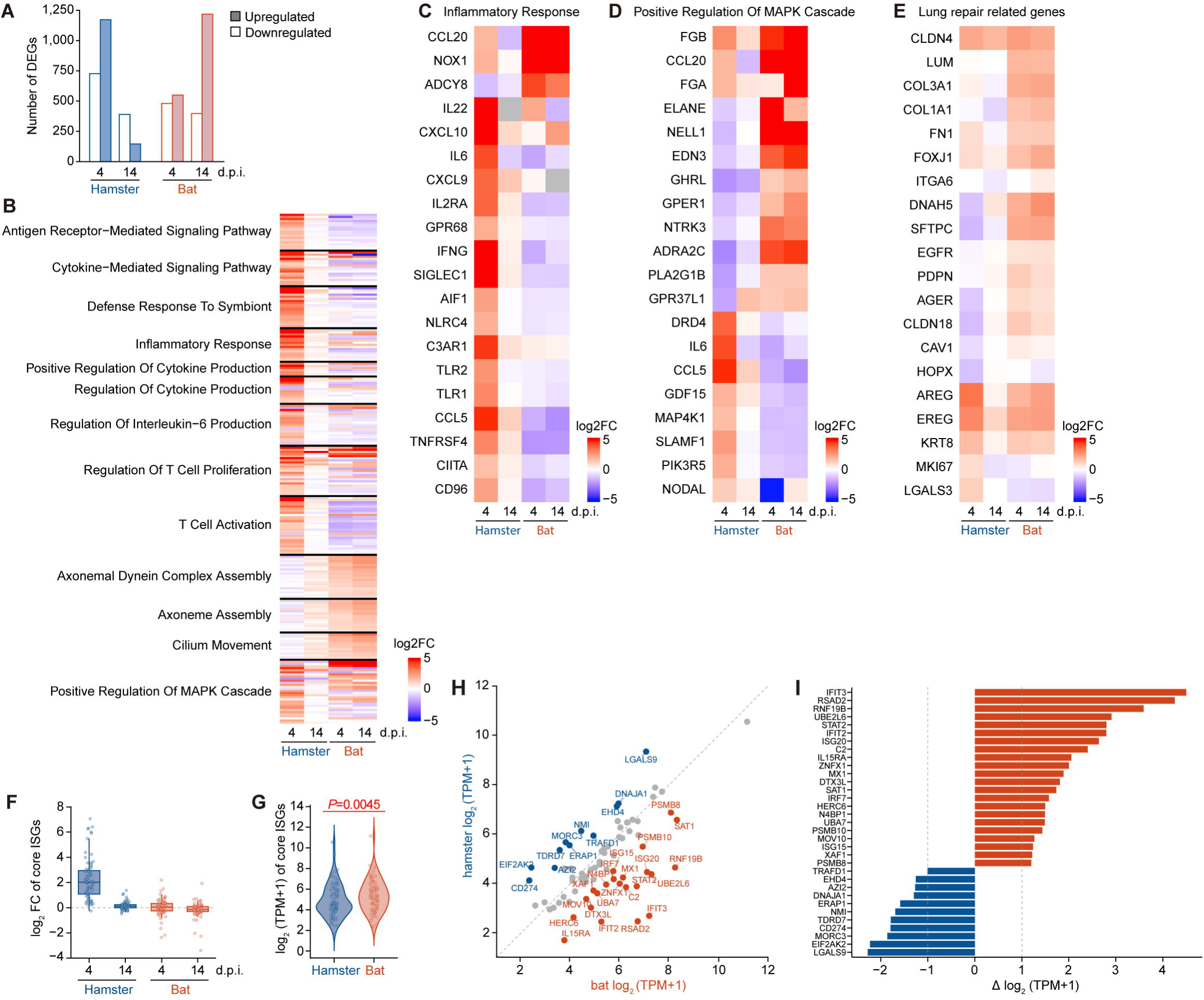
| Distinct transcriptional responses in hamster and bat lungs following SARS-CoV-2 infection. Bulk RNA sequencing was performed on lung tissues collected at 4 and 14 d.p.i. from SARS-CoV-2–infected Syrian hamsters (mock, n = 4; 4 d.p.i., n = 4; 14 d.p.i., n = 4) and greater horseshoe bats (mock, n = 3; 4 d.p.i., n = 3; 14 d.p.i., n = 3). A, Number of DEGs relative to mock controls at 4 and 14 d.p.i. DEGs were defined as genes with Benjamini–Hochberg adjusted P < 0.05 (Wald test) and |log2 fold change (log2FC)| > 1. **B**, Heatmap showing log2FC values of genes belonging to selected GO Biological Process terms identified by enrichment analysis using clusterProfiler. GO enrichment was performed independently for each condition using DEGs as input, and representative terms were selected based on interspecies divergence and redundancy reduction (see Methods). Color scale indicates gene-wise Z-score of log2FC. **C–E**, Heatmaps of representative genes associated with inflammatory response (**C**), positive regulation of MAPK cascade (**D**), and lung repair/regeneration–related processes (**E**). Genes were ranked based on interspecies divergence in log2FC at 4 or 14 d.p.i. Color scale indicates log2FC. **F**, Distribution of log2FC values of conserved ISGs for 4 d.p.i. vs mock and 14 d.p.i. vs mock contrasts in each species. Log2FC values were obtained from differential expression analyses restricted to orthologous genes shared between hamsters and bats, and visualized at the gene level. **G–H**, Baseline expression levels of conserved ISGs at mock (0 d.p.i.), shown as gene-level log2(TPM+1) distributions in boxplot (**G**) and scatterplot (**H**). Baseline expression was defined as the mean across replicate mock samples. Interspecies differences were assessed using a paired Wilcoxon signed-rank test across orthologous genes. **I**, Gene-level baseline expression differences (Δ log2(TPM+1), bat minus hamster) for conserved ISGs. Genes with |Δ| > 1 are highlighted. Conserved ISGs were defined according to the “core mammalian ISG” set described by Shaw et al.^40^ Data are presented as mean ± s.e.m. unless otherwise indicated.

Gene Ontology (GO) enrichment analysis for Biological Process terms was performed on DEGs for four experimental groups. In hamsters at 4 d.p.i., genes annotated with innate immunity and inflammatory response-related GO terms, such as “Cytokine-Mediated Signaling Pathway”, “Defense Response to Symbiont”, and “Inflammatory Response”, exhibited higher log2 fold change (log2FC) score compared to other experimental groups (**Fig. 3B**). In contrast, in bats, the genes annotated in these terms showed lower log2FC score at both 4 and 14 d.p.i. compared to hamsters at 4 d.p.i (**Fig. 3B**). To assess the gene sets related to inflammatory response in bats in detail, the genes annotated with “Inflammatory Response” were compared with log2FC values (**Fig. 3C**). Representative inflammatory mediators, including *IL6*, *IFNG*, and *CXCL9*/*10*, showed marked upregulation at 4 d.p.i. in hamsters, whereas such induction was modest at both 4 and 14 d.p.i. in bats (**Fig. 3C**).

In bats, genes annotated with remodeling cilium-related GO terms, such as “Axoneme Assembly” and “Cilium Movement,” showed higher log2FC score compared to hamsters (**Fig. 3B**).

The GO term “Positive Regulation of MAPK Cascade” showed elevated log2FC score in both species at 4 d.p.i. in hamsters and at both 4 and 14 d.p.i. in bats (**Fig. 3B**). However, gene-level analysis revealed that bats engage this pathway for distinct biological functions from those in hamsters **(Fig. 3D**). In hamsters, inflammatory cytokines and chemokines such as *CCL5* and *GDF15*, were preferentially upregulated (**Fig. 3D**). On the other hand, in bats, genes encoding fibrinogen components, including *FGB* and *FGA* which are potentially associated with tissue remodeling were preferentially upregulated (**Fig. 3D**).

Collectively, these analyses indicated that bats preferentially activate pathways related to tissue repair rather than immune and inflammatory responses upon SARS-CoV-2 infection. To further evaluate this possibility, we analyzed the expression of genes associated with alveolar epithelial repair and regeneration, including markers of alveolar type II (AT2) cells, KRT8+ transitional epithelial cells, which act as critical progenitor cells during lung injury repair, and alveolar type I (AT1) cell differentiation (**Fig. 3E**). In hamsters, a subset of regeneration-associated genes, including the *EGFR* ligands *AREG* and *EREG* and the transitional epithelial cells marker *KRT8*, were transiently upregulated at 4 d.p.i., but their expression declined toward baseline levels by 14 d.p.i. In contrast, bats exhibited broader and more sustained upregulation of regeneration-associated genes at both 4 and 14 d.p.i. Notably, genes associated with AT2 cell activation (*SFTPC*, *EGFR*), transitional epithelial states (*KRT8*, *LGALS3*, *ITGA6*), and AT1 cell differentiation (*AGER*, *PDPN*, *CAV1*, *HOPX*) were consistently upregulated in bats, suggesting activation of an alveolar epithelial regeneration program.

Gene set enrichment analysis (GSEA) further supported the GO enrichment analysis: immune- and inflammatory response–related gene sets were enriched among genes upregulated in hamsters, whereas gene sets associated with tissue repair and regeneration were preferentially enriched among genes upregulated in bats (**Supplementary Fig. 9**).

Our DEG-based analyses suggested that horseshoe bats do not robustly activate innate immune–related pathways upon SARS-CoV-2 infection. On the other hand, previous papers have demonstrated that frugivorous megabats constitutively express some IFN-stimulated genes (ISGs)^13,39^. To address whether constitutive ISG expression is also present in horseshoe bats, we analyzed the expression of mammalian core ISGs conserved among mammals^40^. Comparison of log2 fold changes revealed that substantial ISG induction was observed in hamsters at 4 d.p.i. but not in bats consistent with our enrichment analyses (**Fig. 3C,F**). We next investigated basal ISG expression levels in uninfected group (0 d.p.i.) using transcripts per million (TPM)-normalized values to assess baseline expression states (**Fig. 3G**). Basal ISG expression levels in bats were significantly higher than that in hamsters (1.6-fold median change) (**Fig. 3G**; P = 0.0045). Focusing on individual genes, *MX1* exhibited higher basal ISG expression compared to hamsters (**Fig. 3H,I**), which was consistent with MX1 IHC observations (**Fig. 2J**). Several ISGs demonstrated to play an antiviral function in humans, such as *IFIT2*, *IFIT3*, *ISG20*, *RSAD2*, *MOV10*, and *STAT2*, showed twofold higher TPM values in bats than hamsters.

## Discussion

Our study provides the first investigation of SARS-CoV-2 infection dynamics and host responses in horseshoe bats, natural reservoir hosts of sarbecoviruses, by *in vivo* experimental infection. We found that SARS-CoV-2 replication in horseshoe bats is relatively persistent at a lower level compared to hamsters (**Fig. 1E–H**). Infected horseshoe bats did not exhibit significant body weight loss, suggesting the lack of overt systemic disease upon SARS-CoV-2 infection (**Fig. 1D**). These observations are consistent with the previous reports employing other human-pathogenic virus and natural host bat species, such as Marburg virus infection to Egyptian rousette bats^15,19^.

RNA-seq analysis revealed that horseshoe bats constitutively expressed multiple ISGs, including *MX1*, *IFIT2*, *IFIT3*, *ISG20*, *RSAD2*, *MOV10*, and *STAT2*, in the uninfected group (**Fig. 3G–I**). Such baseline constitutive expression of ISGs is consistent with previous reports in *Pteropus Alecto*, the other bat species^13,39^. Notably, these ISGs are not major inducers of inflammation but directly restrict viral replication in humans at least. For example, MX1 and RSAD2 interfere with viral replication complexes^41–43^, IFIT family proteins inhibit viral RNA translation^44^, and ISG20 degrades viral RNA^45^. In fact, IHC analysis demonstrated constitutive MX1 expression in bronchial epithelial cells (**Fig. 2J**). Taken together, horseshoe bats may maintain a baseline antiviral state in the respiratory epithelium, the primary site of viral entry, by constitutive expression of several ISGs that directly inhibit viral replication. This pre-armed antiviral system may enable horseshoe bats to control sarbecovirus replication at the earliest stage of infection, leading to low-level viral replication and absence of systemic disease.

Neutralization assays showed that humoral immune responses in horseshoe bats were relatively weak compared to hamsters (**Fig. 1I**). Such limited neutralizing antibody responses might explain the persistence of low-level viral RNA. On the other hand, IHC analyses revealed that horseshoe bats exhibited well-organized recruitment of IBA1-positive cells (macrophages) and CD3-positive cells (T cells) to lesion sites in the lung upon SARS-CoV-2 infection (**Fig. 2E, 2F, 2K, and 2L**). This efficient accumulation of macrophages and T cells to infected area may prevent viral dissemination and excessive inflammation throughout the lung. Given the limited understanding of the adaptive immune system in bats, further studies are required to clarify the role of adaptive immunity in viral control of bats.

Our enrichment and GSEA analyses revealed preferential activation of tissue repair–related pathways, rather than innate immunity–related pathways, in SARS-CoV-2–infected horseshoe bats (**Fig. 3B–E and Supplementary Fig. 9**). Lv et al.^46^ have conducted a single-cell transcriptomic analysis of wild *R. affinis* and showed that identified naturally infected lung cells of wild *R. affinis* exhibited upregulation of protein synthesis and tissue repair–related pathways. Altogether, we hypothesized that horseshoe bats appear to prioritize tissue repair and immune regulation over excessive inflammatory responses during viral infection in the lungs. Consistent with this notion, Jayaprakash et al.^17^ have reported that Egyptian rousette bats, the natural hosts of filoviruses, showed activation of tissue repair–related transcriptional programs upon experimental infection of filovirus. This suggests that repair-oriented immune responses may not be specific to horseshoe bats upon sarbecovirus infection^17^. Such a repair-oriented immune strategy may contribute to disease tolerance in bats, enabling viral control without overt tissue damage or clinical disease.

This study has several limitations. First, although all animals were screened prior to infection using pan-sarbecovirus PCR and serological testing against SARS-CoV-2 RBD reactivity, prior exposure to untested viruses cannot be completely excluded and may have influenced baseline immune status. Second, only intranasal inoculation was performed to facilitate comparison with hamster models. However, the natural routes of sarbecovirus transmission in horseshoe bats remain unclear. Future studies using alternative exposure routes may better reflect natural infection conditions.

In summary, this study demonstrates that horseshoe bats mount a distinct host response to SARS-CoV-2 infection, characterized by effective containment of viral replication without the development of systemic disease. Our findings suggest that the antiviral responses in horseshoe bats may involve: (1) a constitutive antiviral state in the respiratory epithelium that suppresses viral replication at the earliest stage of infection; (2) preferential activation of tissue repair–associated pathways; and (3) avoidance of excessive inflammatory and adaptive immune activation. These immune features may enable horseshoe bats to control sarbecovirus replication without developing severe disease, supporting their role as natural reservoir hosts of sarbecoviruses.

## Methods

### Ethics approval

The greater horsehsoe bats (*R. ferrumequinuim*) capturing and collections were conducted by the authors under permission. The bats used in this study were captured in Ishikawa, Niigata and Chiba prefectures, Japan, during May and June 2024–2025 (permission no. 20840–20842 for Ishikawa Prefecture in 2024; permission no. 1251–1256 and 1193–1198 for Niigata Prefecture in 2024 and 2025; and permission no. 336- 1–6 and no. 11-1–6 for Chiba Prefecture in 2024 and 2025) (**Supplementary Table 1**). To minimize the impact on wild populations, only adult males were captured.

For the quarantine of captured bats, anal swabs and plasma samples were collected from captured bats. Anal swabs were screened for sarbecovirus RNA by RT–qPCR using pan-sarbecovirus primers: sarbeco_E_qPCR_F Fw (5’-TCGGAAGAGACAGGTACGTT-3’), sarbeco_E_qPCR_R Rv (5’-TCGAAGCGCAGTAAGGATGG-3’)^34^. Plasma samples were tested for SARS-CoV-2 receptor-binding domain (RBD)-specific antibodies by antigen enzyme-linked immunosorbent assay (ELISA) kit (GenScript, Cat# L00847-A) to confirm seronegativity prior to experimental infection.

### Bat maintenance in ABSL-3 room

Captured bats were maintained in mesh cages placed inside negative-pressure isolators within an ABSL-3 facility in The Institute of Medical Science, The University of Tokyo (IMSUT), Japan. Each cage housed three bats throughout the study. When we conduct experimental procedures, the closed isolators were moved to a safety cabinet in ABSL-3 facility, and the bats were handled in the safety cabinet. The live bats were handled in ABSL-3 facility, with personnel wearing full-body personal protective equipment and N95 masks. All experimental facilities have been approved by IMSUT.

### Bat education for self-feeding in ABSL-3 room

Bats were gradually acclimated to self-feeding on mealworms through a stepwise training protocol: (i) direct feeding using forceps; (ii) placing mealworms in a tray and gradually bringing the tray closer to the bats to train them to feed directly from the tray; and (iii) independent feeding from trays placed inside the cage (**Fig. 1B** and **Supplementary Movie S1**). This acclimation process typically required 2–4 weeks (**Supplementary Fig. 1**). The ABSL-3 facility was maintained at approximately 21°C, and humidity was controlled by periodic misting of the enclosures.

### Cell culture

Lenti-X 293T cells (Takara, Cat# 632180) and HOS cells (human osteosarcoma cell line; ATCC CRL-1543) engineered to stably express human ACE2 and TMPRSS2, hamster (*M. auratus*) ACE2 together with human TMPRSS2, or bat (*R. ferrumequinum*) ACE2 together with human TMPRSS2 were used in this study. All cells were cultured in high-glucose Dulbecco’s modified Eagle’s medium (DMEM; Wako, Cat# 044-29765) supplemented with 10% fetal bovine serum (Sigma-Aldrich, Cat# 172012-500ML), and 100 units penicillin, 100 µg/ml streptomycin (Sigma-Aldrich, Cat# P4333-100ML), zeocin (50 μg/mL; InvivoGen, Cat# ant-zn-1), and G418 (400 μg/mL; Nacalai Tesque, Cat# G8168-10ML). VeroE6/TMPRSS2 cells (VeroE6 cells stably expressing human TMPRSS2; JCRB Cell Bank, JCRB1819) (PMID: 32165541) were maintained in DMEM (low glucose) (Wako, Cat# 041-29775) containing 10% FBS, G418 (1 mg/ml; Nacalai Tesque, Cat# G8168-10ML) and 100 units penicillin, 100 µg/ml streptomycin (Sigma-Aldrich, Cat# P4333-100ML).

### Plasmid construction

Plasmids encoding human codon-optimized S-proteins of selected sarbecoviruses were prepared in our previous study^33,47,48^. Codon-optimized S expression plasmids for SARS-CoV-2 Wuhan and Delta S:Q493K were generated by site-directed mutagenesis PCR by using following primers: pC-S_SARS-CoV-2_Q493K_Fw Fw (5’-TGTTACTTTCCACTCAAGTCCTATGGCTTCCAA-3’), pC-S_SARS-CoV-2_Q493K _Rv Rv (5’-TTGGAAGCCATAGGACTTGAGTGGAAAGTAACA-3’). The amplified fragments were inserted into the pCAGGS expression vector at the KpnI and NotI restriction sites using the In-Fusion HD Cloning Kit (Takara, Cat# Z9650N). All constructs were verified by Sanger sequencing (Eurofins), and sequence analyses were performed using SnapGene v8.0.1 (SnapGene software).

### Pseudovirus assay

Pseudovirus assay was performed as previously described^49–53^. HIV-1–based, luciferase-encoding reporter viruses were pseudotyped with S proteins of sarbecoviruses and their derivatives. Lenti-293T cells (500,000 cells) were cotransfected with 0.8 μg psPAX2-IN/HiBiT^48^, 0.8 μg pWPI-Luc2^54^, and 0.4 μg plasmids encoding the parental S protein or its derivatives using TransIT-293 (Takara, Cat# MIR2704) according to the manufacturer’s instructions. Culture supernatants containing pseudoviruses were collected at 48 h posttransfection and stored at –80 °C until use. For infection assays, viral input was normalized based on HiBiT luminescence quantified using the Nano-Glo HiBiT Lytic detection system (Promega, Cat# N3040) as previously described^55^. Target cells consisted of HOS-TMPRSS2 cells stably expressing human and bat (*R. ferrumequinum*) ACE2. At 48 h postinfection, infected cells were lysed with Bright-Glo Luciferase Assay System (Promega, Cat# E2620), and luminescence was measured using a GloMax Explorer multimode microplate reader (Promega).

### SARS-CoV-2 reverse genetics

Recombinant SARS-CoV-2 Delta and Delta S:Q493K was generated using a circular polymerase extension reaction (CPER) as previously described^31,52,56^ with modifications^57^. In brief, nine overlapping DNA fragments covering the SARS-CoV-2 genome were amplified by PCR using PrimeSTAR GXL DNA polymerase (Takara, Cat# R050A). A linker fragment encoding hepatitis delta virus ribozyme, bovine growth hormone poly A signal and cytomegalovirus promoter was also prepared by PCR. These ten fragments were combined and subjected to CPER to assemble the full-length viral genome^56^. For generation of the Delta S:Q493K variant, fragment 8 encoding the Spike region was modified to introduce the Q493K substitution. The mutated fragment 8 was constructed by site-directed overlap extension PCR using the following primers: pCSII_F8_Q493K Fw (5’- TGTTACTTTCCTTTAAAATCATATGGTTTCCAA-3’), pCSII_F8_Q493K Rv (5’- TTGGAAACCATATGATTTTAAAGGAAAGTAACA-3’). Nucleotide sequences were determined by Sanger sequencing (Fasmac), and the sequence data were analyzed by SnapGene v8.0.1 (SnapGene software).

To produce infectious recombinant virus, the CPER products (25 µl) were transfected into VeroE6/TMPRSS2 cells with TransIT-X2 (Takara, Cat# MIR6003) according to the manufacturer’s protocol. When CPE was observed, supernatants were harvested. To eliminate residual CPER-derived DNA (i.e., SARS-CoV-2-related DNA), 1 ml of the seed virus was treated with 2 μl TURBO DNase (Thermo Fisher Scientific, Cat# AM2238) at 37°C for 1 h. Complete removal of the CPER products from the seed virus was confirmed by PCR analysis. All the viruses were kept at −80°C until use. The working virus stock was prepared using the seed virus as described below.

### Recombinant SARS-CoV-2 preparation and titration

The working virus stocks of SARS-CoV-2 Delta and Delta S: Q493K were prepared and titrated as previously described^31,58^. In brief, seed virus was inoculated into VeroE6/TMPRSS2 cells (5,000,000 cells in a T-75 flask). At 1 hour post infection, the culture medium was replaced with DMEM (low glucose) (Sigma-Aldrich, Cat# D6046) containing 2% FBS and 1% PS. At 3 d.p.i., the culture medium was harvested and centrifuged, and the supernatants were collected as the working virus stock.

The titer of the prepared working virus was measured as the 50% tissue culture infectious dose (TCID50). Briefly, 1 day before infection, VeroE6/TMPRSS2 (10,000 cells) were seeded into a 96-well plate. Serially diluted virus stocks were inoculated into the cells and incubated at 37°C for 4 days. The cells were observed under a microscope to judge the CPE appearance. The value of TCID50/ml was calculated with the Reed–Muench method^59^. For verification of the sequences of working viruses, viral RNA was extracted from the working viruses using a QIAamp viral RNA mini kit (Qiagen, Cat# 52906) and viral genome sequences were analyzed as described (see “Viral genome sequencing” section).

### Live virus genome sequencing

Viral genome sequencing was performed as previously described^37^. Briefly, the virus sequences were verified by viral RNA-sequencing analysis. Viral RNA was extracted using a QIAamp viral RNA mini kit (Qiagen, Cat# 52906). The sequencing library employed for total RNA sequencing was prepared using the NEBNext Ultra RNA Library Prep Kit for Illumina (New England Biolabs, Cat# E7530). Paired-end 76-bp sequencing was performed using a MiSeq system (Illumina) with MiSeq reagent kit v3 (Illumina, Cat# MS-102-3001). Sequencing reads were trimmed using fastp v0.21.0^60^ and subsequently mapped to the viral genome sequences of a lineage A isolate (strain WK-521, GISAID ID: EPI_ISL_408667)^61^ using BWA-MEM v0.7.17^62^. Variant calling, filtering, and annotation were performed using SAMtools v1.9^63^ and snpEff v5.0e^64^. Information on the unexpected substitutions detected is summarized in Supplementary Table 2.

### Virus infection in cells

HOS-T cells stably expressing human ACE2, hamster (*M. auratus*) ACE2, or bat (*R. ferrumequinum*) ACE2 were seeded in 96-well plates at a density of 20,000 cells per well one day prior to infection. Cells were infected with SARS-CoV-2 Delta Q493K at a multiplicity of infection (MOI) of 0.1 and incubated at 37 °C for 1 h to allow viral adsorption. The infected cells were washed, and 180 μl of culture medium was added. At the indicated time points post-infection, 10 μl of culture supernatant was collected for quantification of viral RNA by RT–qPCR as described below.

### RT-qPCR

RT–qPCR was conducted as previously described^31,52^. In brief, 5 μl of culture supernatant was mixed with 5 μl of 2× RNA lysis buffer (2% Triton X-100, 50 mM KCl, 100 mM Tris-HCl (pH 7.4), 40% glycerol and 0.8 U/μl recombinant RNase inhibitor (Takara, 2313B)) and incubated at room temperature for 10 min. RNase-free water (90 μl) was added, and the diluted sample (2.5 μl) was used as the template for real-time RT–PCR performed according to the manufacturer’s protocol using the One Step TB Green PrimeScript PLUS RT-PCR kit (Takara, RR096A) and the following primers^31^: Forward_N Fw (5’-AGCCTCTTCTCGTTCCTCATCAC-3’), Reverse_N Rv (5’-CCGCCATTGCCAGCCATTC-3’).

### Experimental infection in animals and sample collection

Syrian hamsters (male, 4 weeks old) were purchased from Japan SLC and divided into groups by simple randomization. Baseline body weights were measured before infection for both bats and hamsters. For the virus infection experiments, we anaesthetized bats and hamsters by intramuscular injection of a mixture of 0.15 mg/kg medetomidine hydrochloride (Domitor, Nippon Zenyaku Kogyo), 2.0 mg/kg midazolam (Dormicum, FUJIFILM Wako Chemicals) and 2.5 mg/kg butorphanol (Vetorphale, Meiji Seika Pharma). The Delta S:Q493K (100,000 TCID50 in 75 μl) or saline (75 μl) were intranasally inoculated under anesthesia. Oral and anal swabs were daily collected until 21 d.p.i., and every 3 days after that to 28 d.p.i (**Fig. 1C**). Body weight was measured for same frequency to swabs (**Fig. 1C**). Some individuals were sacrificed by cervical dislocation at 0, 4, 7, 14, 28 d.p.i. and trachea, lung, stomach, and intestine were anatomically collected. These tissues were used for histopathological analyses as below. Only lung of these tissues was also used for RNA-seq analysis. Viral RNA load in the oral, anal swabs and lung was determined by RT–qPCR. RT–PCR performed according to the manufacturer’s protocol using the One Step TB Green PrimeScript PLUS RT-PCR kit (Takara, RR096A) and the following primers^31^: Forward_N Fw (5’-AGCCTCTTCTCGTTCCTCATCAC-3’), Reverse_N Rv (5’-CCGCCATTGCCAGCCATTC-3’). Viral titers in the lung were determined by TCID50. Plasmas were collected at 0, 14, 28 d.p.i. for neutralization assay as below. No method of randomization was used to determine how the animals were allocated to the experimental groups and processed in this study because covariates (sex and age) were identical for hamsters and no way to know age for bats. The number of investigators was limited, as most of experiments were performed in high-containment laboratories. Therefore, blinding was not carried out.

### Neutralization assays

Neutralization assays were performed as previously described^38,53,65^. For human sera, the SARS-CoV-2 Delta and Delta S:Q493K pseudoviruses (counting ∼100,000 relative light units) were incubated with serially diluted (120-fold to 87,480-fold dilution at the final concentration) heat-inactivated 3-dose vaccinated sera of human at 37°C for 1 hour.

For hamsters and bats plasma, the SARS-CoV-2 Delta and Delta S:Q493K pseudoviruses (counting ∼100,000 relative light units) were incubated with serially diluted (80-fold to 58,320-fold dilution at the final concentration) heat-inactivated plasma of hamsters and bats at 37°C for 1 hour. Pseudoviruses without sera were included as controls. Then, 20 μl mixture of pseudovirus and serum or plasma was added to HOS-TMPRSS2 human ACE2 cells (10,000 cells/100 μl) in a 96-well white plate. Two days post infection, the infected cells were lysed with a Bright-Glo luciferase assay system (Promega, Cat# E2620), and the luminescent signal was measured using a GloMax explorer multimode microplate reader 3500 (Promega). The assay of each serum sample was performed in triplicate, and the 50% neutralization titer (NT50) was calculated using Prism 9 (GraphPad Software).

### H&E staining

Briefly, excised animal tissues were fixed with 10% formalin neutral buffer solution and processed for paraffin embedding. The paraffin blocks were sectioned at a thickness of 3 μm and then mounted on MAS-GPcoated glass slides (Matsunami Glass, Cat# S9901). The deparaffinized sections were stained with H&E following previously described protocol^31^ (**Fig. 2**).

### Immunohistochemistry

Immunohistochemical (IHC) analysis (**Fig. 2**) was performed using an Autostainer Link 48 (Dako). The deparaffinized sections were exposed to EnVision FLEX target retrieval solution high pH (Agilent, Cat# K8004) for 20 min at 97 ◦C for activation, and a mouse anti-SARS-CoV-2 N monoclonal antibody (clone 1035111, R&D Systems, Cat# MAB10474-SP, 1:400) was used as a primary antibody. For only lung tissue, IHC using anti-MX1 polyclonal antibody (Proteintech, Cat# 13750-1-AP, 1:200), anti-Iba-1 polyclonal antibody (FUJIFILM Wako, Cat# 019-19741, 1:1,000), and anti-CD3 polyclonal antibody (DAKO, Cat# IR503, RTU) were also performed. The sections were sensitized using EnVision FLEX for 15 min and visualized by peroxidase-based enzymatic reaction with 3,3′ -diaminobenzidine tetrahydrochloride (Dako, Cat# DM827) as substrate for 5 min. The IHC staining positivity was evaluated by NDP.scan software v3.2.4 (Hamamatsu Photonics). The IHC staining positivity was measured as the area and calculated as the percentage using Fiji software v2.2.0 (ImageJ).

### RNA extraction and RNA-seq

Total RNA was extracted from the lung parenchyma using QIAamp RNA Blood Mini Kit (Qiagen, Cat# 52304) following the manufacturer’s protocols. cDNA library was subsequently synthesized using TruSeq Stranded mRNA Library Prep (96 Samples) (Illumina, Cat# 20020595) and IDT for Illumina – TruSeq RNA UD Indexes v2 (96 Indexes, 96 Samples) (Illumina, Cat# 20040871) following the manufacturer’s protocols. The library was then sequenced using the AVITI system (Element Biosciences). Nucleotide sequences in all samples have a mean quality score ≥35.94, ensuring the overall accuracy of the base calling > 99.97% (**Supplementary Table 3**). Samples were confirmed with RNA integrity number higher that 7.5 by Bioanalyzer (Agilent) (**Supplementary Table 3**).

### Transcriptomic read pre-processing and quantification

Transcriptomic reads were pre-processed to ensure the quality of read alignment. First, adapter trimming and read filtering were performed using fastp v0.23.4 in the default mode^60^. Then, the trimmed reads were mapped to the Syrian hamster (GCF_017639785.1), horseshoe bat (GCF_004115265.2) genomes using STAR aligner v 2.7.11a ^66^ with 5, 500,000, 1,000,000, 1, 0.33, and 0.33 as the maximum number of mismatches per pair, maximum intron length, maximum gaps between two mates, minimum overhang for annotated spliced alignments, minimum number of bases matched over read length, and minimum alignment score over read length, respectively, different from the default mode. All read samples contain ≥70.31% of mapped reads, indicating the lack of substantial contamination from other organisms (**Supplementary Table 3**). After that, the mapped reads were quantified using featureCounts provided in the subread package v.2.0.6 with 0.25 as the minimum fraction of overlapping bases^67^.

### DEG analysis

Differential expression and functional enrichment analyses were performed using R v4.3.2^68^. Differential expression analysis was then performed using DEseq2 v1.42.1^69^ with log2 fold change shrinkage estimated using the apeglm method. For the hamster dataset. The samples sacrificed at 0 d.p.i. were used as control for the infected samples at both 4 and 14 d.p.i. In the pairwise comparison. In pairwise comparisons, DEGs were defined as those with a Benjamini–Hochberg adjusted P value < 0.05 (Wald test) and an absolute log2 fold change (|log2FC|) greater than 1. The hamster-greater horseshoe bats orthologous relationship data were collected from the NCBI Gene database^70^ on October 1, 2025 for comparing the results between species.

### GO enrichment analysis

Gene Ontology (GO) enrichment analysis was performed independently for each experimental condition using the clusterProfiler R package (v4.10.1)^71^, focusing on GO Biological Process terms. DEGs were used as input, and terms with Benjamini–Hochberg adjusted P < 0.05 were considered significant. For each condition, the top 10 terms ranked by adjusted P value were extracted, and their union across bat 4 d.p.i. (Bat D4), bat 14 d.p.i. (Bat D14), hamster 4 d.p.i. (Ham D4), and hamster 14 d.p.i. (Ham D14) was defined as the candidate GO term set (n = 40). Enrichment strength was quantified as −log10(adjusted P value). A term-level divergence score was defined as the absolute difference between Ham D4 and the maximum enrichment observed in bats (Bat D4 or Bat D14), and the top 15 divergent terms were selected. Redundancy was reduced by calculating pairwise Jaccard similarity coefficients based on overlapping DEG sets; for term pairs with similarity ≥ 0.6, only the higher-scoring term was retained. For selected GO terms, gene-level log2FC values across Bat D4, Bat D14, Ham D4, and Ham D14 were visualized as heatmaps (**Fig. 3B**).

For selected GO terms, gene-level expression patterns were further examined. In addition, a custom lung repair/regeneration gene set was curated based on established markers of AT2/AT1 epithelial cells, KRT8⁺ transitional cells, and extracellular matrix remodeling. A gene-level divergence score was defined as the maximum absolute difference in log2FC between bats and hamsters at D4 or D14. Genes were ranked by this score, and the top 20 genes per term were visualized. Heatmaps were generated using log2FC values across the four conditions.

### GSEA

GSEA was performed using the clusterProfiler R package (v4.10.1). For each species (bat and hamster) and each comparison (0 vs 4 d.p.i. and 0 vs 14 d.p.i.), a ranked gene list was generated from differential expression results using log2FC, restricted to ortholog-mapped genes (“orthologsOnly”). Gene symbols were converted to Entrez Gene IDs using bitr (clusterProfiler) with the org.Hs.eg.db annotation package, and genes without Entrez ID mapping were excluded. The ranked list was sorted in decreasing order of log2FC. GSEA was conducted against Gene Ontology Biological Process (GO BP) gene sets using gseGO (ont = “BP”, minGSSize = 10, pvalueCutoff = 0.05). Adjusted P values were calculated using the Benjamini–Hochberg method. For visualization, results were converted to a data frame, entries with missing adjusted P values were excluded, and the top 20 GO BP terms ranked by adjusted P value were displayed. Dotplots were generated with normalized enrichment score (NES) on the x-axis, term description on the y-axis, point size proportional to gene ratio (setSize / length(geneList)), and point color representing adjusted P value (**Supplementary Fig. 9**).

### ISG expression analysis

Expression patterns of conserved ISGs were assessed using the “core mammalian ISG” set defined by Shaw et al.^40^. Ortholog-mapped lung RNA-seq differential expression results were used for bats and hamsters. For each species, log2FC values from the D0 vs D4 and D0 vs D14 contrasts were extracted and mapped to human ortholog gene symbols, and core ISGs were selected by symbol matching. ISG log2FC distributions were visualized as boxplots with overlaid per-gene points for each contrast.

Baseline ISG expression was quantified using the same core mammalian ISG set as above. Gene-level read counts from lung RNA-seq were converted to TPM using gene length normalization (RPK) followed by library-size scaling, and values were log2-transformed as log2(TPM+1). Baseline (0 d.p.i.) expression was defined using mock-inoculated samples, including additional mock controls obtained from a previous ligand-administration experiment performed under the same experimental and sequencing workflow, to increase the number of 0 d.p.i. replicates.For each gene, baseline expression (0 d.p.i.) was defined as the mean log2(TPM+1) across replicate samples. Baseline ISG expression distributions were visualized using violin plots, and interspecies differences were assessed across genes using a paired Wilcoxon signed-rank test. Gene-level baseline differences were additionally summarized as Δlog2(TPM+1) (bat minus hamster), and genes with |Δ| > 1 were highlighted in bar and scatter plots.

## Data and code availability

All transcriptomic data from bat and hamster lungs are available from Gene Expression Omnibus repository (GEOXXXX). Code and data used in this study are available on the following GitHub repository: https://github.com/TheSatoLab/XXXXXXXXX.

## Acknowledgements

We thank all members of Division of Systems Virology, especially Naoko Misawa, Ziyi Guo, and Chen Luo for their assistance with bat maintenance; Shusuke Kawakubo, Wenye Li, Arnon Plianchaisuk and Jarel Elgin Tolentino for their assistance with bat sampling; and Maximilian Stanley Yo for their assistance with bat sampling and proofreading our manuscript.

## Supplementary Figures

**Supplementary Figure 1 | Percentage body weight change before experimental infection.**

Greater horseshoe bats captured from wild caves were trained to acclimate self-feeding on the mealworms prior to experimental infection. **A–E**, Percentage body weight change from the date of capturing to the date of viral inoculation. Since bats were captured on six separate occasions, the acclimation training was conducted in six batches. Vertical dotted lines and specimen IDs indicate the timepoint at which each bat completed acclimation and began self-feeding on mealworms.

**Supplementary Figure 2 | Q493K mutation in the spike protein enables efficient entry via greater horseshoe bat ACE2.**

**A**, Representative western blot showing expression of FLAG-tagged ACE2 from greater horseshoe bat or Syrian hamster in HOS-T cells. TUBA served as a loading control. **B**, Infectivity assay of pseudovirus bearing spike protein of several SARS-CoV-2 variants in HOS-T cells expressing bat ACE2. Pseudoviruses equivalent to 3,500 pg of HIV-1 p24 per well were used for infection. The Q493K mutation increased the infectivity of both SARS-CoV-2 Delta spike-bearin pseudovirus against bat ACE2. **C**, Growth kinetics of recombinant SARS-CoV-2 Delta virus carrying the Q493K mutation in HOS-T cells expressing bat ACE2. **D**, Growth kinetics of recombinant SARS-CoV-2 Delta virus carrying the Q493K mutation in HOS-T cells expressing hamster ACE2. HOS-T cells were seeded at 20,000 cells per well and infected at a multiplicity of infection (MOI) of 0.1 (**C–D**). Viral RNA in the supernatant was quantified by RT–qPCR (**C–D**). This assay demonstrated that the Q493K mutation increased the efficiency of bat ACE2 usage even in live viruses. On the other hand, this mutation did not affect efficiency of hamster ACE2 usage of Delta, though these viruses can substantially replicate in these cells.

**Supplementary Figure 3 | Q493K mutation in the spike protein does not affect efficiency of entry via human ACE2.**

**A**, Pseudovirus infectivity assay using HOS-T cells expressing human ACE2. The Q493K mutation did not affect the infectivity of both SARS-CoV-2 Wuhan and Delta spike pseudoviruses in HOS-T cells expressing human ACE2. **B**, Growth kinetics of recombinant SARS-CoV-2 Delta virus carrying the Q493K mutation in HOS-T cells expressing human ACE2. The HOS-T cells were seeded at 20,000 cells per well and infected at a multiplicity of infection (MOI) of 0.1. Viral RNA in the culture supernatant was quantified by RT–qPCR. **C**, Neutralizing activity of 3-dose vaccinated sera of human (n=12) against SARS-CoV-2 Delta Q493K spike-bearing pseudotyped virus, determined by pseudovirus neutralization assay.

**Supplementary Figure 4 | Inflammatory lesion area in the lungs of infected hamsters and bats.**

**A**, Lung of the hamsters infected with Delta carrying Q493K mutation stained with H&E. **B**, Lung of the greater horseshoe bats infected with Delta carrying Q493K mutation stained with H&E. In each timepoint, H&E staining (top) and the digitalized inflammation area (bottom, indicated in red) are shown. The number in the panel of the digitalized inflammation area indicates the percentage of the section represented by the indicated area (that is, the area indicated with red color per the total area of the lung). Scale bars, 5 mm.

**Supplementary Figure 5 | N-protein-positive area in the lungs of infected hamsters and bats.**

**A**, IHC for SARS-CoV-2 N protein in the lung of the hamsters infected with Delta carrying Q493K mutation. **B**, IHC for SARS-CoV-2 N protein in the lung of the greater horseshoe bats infected with Delta carrying Q493K mutation. In each timepoint, IHC (top) and the digitalized N-positive area (bottom, indicated in red) are shown. The number in the panel of the digitalized N-positive area indicates the percentage of the section represented by the indicated area (that is, the area indicated with red color per the total area of the lung). Scale bars, 5 mm.

**Supplementary Figure 6 | MX1-positive area in the lungs of infected hamsters and bats.**

**A**, IHC for MX1 protein in the lung of the hamsters infected with Delta carrying Q493K mutation. **B**, IHC for MX1 protein in the lung of the greater horseshoe bats infected with Delta carrying Q493K mutation. In each timepoint, IHC (top) and the digitalized MX1-positive area (bottom, indicated in red) are shown. The number in the panel of the digitalized MX1 -positive area indicates the percentage of the section represented by the indicated area (that is, the area indicated with red color per the total area of the lung). Scale bars, 5 mm.

**Supplementary Figure 7 | IBA1-positive area in the lungs of infected hamsters and bats.**

**A**, IHC for IBA1 protein in the lung of the hamsters infected with Delta carrying Q493K mutation. **B**, IHC for IBA1 protein in the lung of the greater horseshoe bats infected with Delta carrying Q493K mutation. In each timepoint, IHC (top) and the digitalized IBA1-positive area (bottom, indicated in red) are shown. The number in the panel of the digitalized IBA1-positive area indicates the percentage of the section represented by the indicated area (that is, the area indicated with red color per the total area of the lung). Scale bars, 5 mm.

**Supplementary Figure 8 | CD3-positive area in the lungs of infected hamsters and bats.**

**A**, IHC for CD3 protein in the lung of the hamsters infected with Delta carrying Q493K mutation. **B**, IHC for CD3 protein in the lung of the greater horseshoe bats infected with Delta carrying Q493K mutation. In each timepoint, IHC (top) and the digitalized CD3-positive area (bottom, indicated in red) are shown. The number in the panel of the digitalized CD3-positive area indicates the percentage of the section represented by the indicated area (that is, the area indicated with red color per the total area of the lung). Scale bars, 5 mm.

**Supplementary Figure 9 | GSEA of lung transcriptomes following SARS-CoV-2 infection in hamsters and greater horseshoe bats.**

GSEA was performed using GO Biological Process terms based on ranked gene lists derived from differential expression analyses. The top 20 enriched pathways are shown. Dot size indicates the GeneRatio and color represents the adjusted P value (Benjamini–Hochberg correction). The x-axis shows the NES. **A**, Enriched GO terms in hamster lungs at 4 d.p.i. **B**, Enriched GO terms in hamster lungs at 14 d.p.i. **C**, Enriched GO terms in greater horseshoe bat lungs at 4 d.p.i. **D**, Enriched GO terms in greater horseshoe bat lungs at 14 d.p.i.

## Supplementary Tables

**Supplementary Table 1 | Information of bat specimens regarding captured site, experimental usage.**

**Supplementary Table 2 | Summary of unexpected amino acid mutations detected in the working virus stocks.**

**Supplementary Table 3 | Information of animal samples used in RNA transcriptomic analyses regarding quality of RNA samples, transcriptome reads, and read mapping.**

## Reference

1. Ciotti, M. et al. The COVID-19 pandemic. Critical Reviews in Clinical Laboratory Sciences 57, 365–388 (2020).

2. Hu, B. et al. Discovery of a rich gene pool of bat SARS-related coronaviruses provides new insights into the origin of SARS coronavirus. PLOS Pathogens 13, e1006698 (2017).

3. Worobey, M. et al. The Huanan Seafood Wholesale Market in Wuhan was the early epicenter of the COVID-19 pandemic. Science 377, 951–959 (2022).

4. Boni, M. F. et al. Evolutionary origins of the SARS-CoV-2 sarbecovirus lineage responsible for the COVID-19 pandemic. Nat Microbiol 5, 1408–1417 (2020).

5. Delaune, D. et al. A novel SARS-CoV-2 related coronavirus in bats from Cambodia. Nat Commun 12, 6563 (2021).

6. Li, W. et al. Bats Are Natural Reservoirs of SARS-Like Coronaviruses. Science 310, 676–679 (2005).

7. Ruiz-Aravena, M. et al. Ecology, evolution and spillover of coronaviruses from bats. Nat Rev Microbiol 20, 299–314 (2022).

8. Wacharapluesadee, S. et al. Evidence for SARS-CoV-2 related coronaviruses circulating in bats and pangolins in Southeast Asia. Nat Commun 12, 972 (2021).

9. Zhou, P. et al. A pneumonia outbreak associated with a new coronavirus of probable bat origin. Nature 579, 270–273 (2020).

10. Zhou, H. et al. Identification of novel bat coronaviruses sheds light on the evolutionary origins of SARS-CoV-2 and related viruses. Cell 184, 4380–4391.e14 (2021).

11. Calisher, C. H., Childs, J. E., Field, H. E., Holmes, K. V. & Schountz, T. Bats: Important Reservoir Hosts of Emerging Viruses. Clinical Microbiology Reviews 19, 531–545 (2006).

12. Letko, M., Seifert, S. N., Olival, K. J., Plowright, R. K. & Munster, V. J. Bat-borne virus diversity, spillover and emergence. Nat Rev Microbiol 18, 461–471 (2020).

13. Irving, A. T., Ahn, M., Goh, G., Anderson, D. E. & Wang, L.-F. Lessons from the host defences of bats, a unique viral reservoir. Nature 589, 363–370 (2021).

14. Pei, G., Balkema-Buschmann, A. & Dorhoi, A. Disease tolerance as immune defense strategy in bats: One size fits all? PLOS Pathogens 20, e1012471 (2024).

15. Guito, J. C. et al. Asymptomatic Infection of Marburg Virus Reservoir Bats Is Explained by a Strategy of Immunoprotective Disease Tolerance. Current Biology 31, 257–270.e5 (2021).

16. Halpin, K. et al. Pteropid Bats are Confirmed as the Reservoir Hosts of Henipaviruses: A Comprehensive Experimental Study of Virus Transmission. Am J Trop Med Hyg 85, 946–951 (2011).

17. Jayaprakash, A. D. et al. Marburg and Ebola Virus Infections Elicit a Complex, Muted Inflammatory State in Bats. Viruses 15, (2023).

18. Kessler, S. et al. Jamaican fruit bats mount a stable and highly neutralizing antibody response after bat influenza virus infection. Proceedings of the National Academy of Sciences 121, e2413619121 (2024).

19. Middleton, D. J. et al. Experimental Nipah Virus Infection in Pteropid Bats (*Pteropus poliocephalus*). Journal of Comparative Pathology 136, 266–272 (2007).

20. Paweska, J. T. et al. Virological and Serological Findings in Rousettus aegyptiacus Experimentally Inoculated with Vero Cells-Adapted Hogan Strain of Marburg Virus. PLOS ONE 7, e45479 (2012).

21. Paweska, J. T. et al. Lack of Marburg Virus Transmission From Experimentally Infected to Susceptible In-Contact Egyptian Fruit Bats. J Infect Dis 212, S109–S118 (2015).

22. Schuh, A. J. et al. Modelling filovirus maintenance in nature by experimental transmission of Marburg virus between Egyptian rousette bats. Nat Commun 8, 14446 (2017).

23. Storm, N., Vuren, P. J. V., Markotter, W. & Paweska, J. T. Antibody Responses to Marburg Virus in Egyptian Rousette Bats and Their Role in Protection against Infection. Viruses 10, (2018).

24. Williamson, M. et al. Transmission studies of Hendra virus (equine morbilli-virus) in fruit bats, horses and cats. Australian Veterinary Journal 76, 813–818 (1998).

25. Burke, B. et al. Regulatory T cell-like response to SARS-CoV-2 in Jamaican fruit bats (Artibeus jamaicensis) transduced with human ACE2. PLOS Pathogens 19, e1011728 (2023).

26. Hall, J. S. et al. Experimental Infection of Mexican Free-Tailed Bats (Tadarida brasiliensis) with SARS-CoV-2. mSphere 8, e00263–22 (2023).

27. Port, J. R. et al. Jamaican fruit bat (Artibeus jamaicensis) insusceptibility to mucosal inoculation with SARS-CoV-2 Delta variant is not caused by receptor compatibility. npj Viruses 2, 26 (2024).

28. Schlottau, K. et al. SARS-CoV-2 in fruit bats, ferrets, pigs, and chickens: an experimental transmission study. The Lancet Microbe 1, e218–e225 (2020).

29. Sato, Y. & Sekijima, T. Establishment of a feeding method to promote self-feeding in lesser horseshoe bats under captive conditions. Honyurui Kagaku 59, 3–13 (2019) [in Japanese].

30. Imai, M. et al. Syrian hamsters as a small animal model for SARS-CoV-2 infection and countermeasure development. Proceedings of the National Academy of Sciences 117, 16587–16595 (2020).

31. Saito, A. et al. Enhanced fusogenicity and pathogenicity of SARS-CoV-2 Delta P681R mutation. Nature 602, 300–306 (2022).

32. Sia, S. F. et al. Pathogenesis and transmission of SARS-CoV-2 in golden hamsters. Nature 583, 834–838 (2020).

33. Fujita, S. et al. Determination of the factors responsible for the tropism of SARS-CoV-2-related bat coronaviruses to Rhinolophus bat ACE2. Journal of Virology 97, e00990–23 (2023).

34. Murakami, S. et al. Isolation of Bat Sarbecoviruses, Japan. Emerg Infect Dis 28, 2500–2503 (2022).

35. Fujita, S. et al. Virological characteristics of a SARS-CoV-2-related bat coronavirus, BANAL-20-236. eBioMedicine 104, (2024).

36. Suzuki, R. et al. Attenuated fusogenicity and pathogenicity of SARS-CoV-2 Omicron variant. Nature 603, 700–705 (2022).

37. Yamasoba, D. et al. Virological characteristics of the SARS-CoV-2 Omicron BA.2 spike. Cell 185, 2103–2115.e19 (2022).

38. Yamasoba, D. et al. Virological characteristics of the SARS-CoV-2 omicron XBB.1.16 variant. The Lancet Infectious Diseases 23, 655–656 (2023).

39. Zhou, P. et al. Contraction of the type I IFN locus and unusual constitutive expression of IFN-α in bats. Proceedings of the National Academy of Sciences 113, 2696–2701 (2016).

40. Shaw, A. E. et al. Fundamental properties of the mammalian innate immune system revealed by multispecies comparison of type I interferon responses. PLOS Biology 15, e2004086 (2017).

41. Bizzotto, J. et al. SARS-CoV-2 Infection Boosts *MX1* Antiviral Effector in COVID-19 Patients. iScience 23, 101585 (2020).

42. Haller, O., Kochs, G. & Weber, F. Interferon, Mx, and viral countermeasures. Cytokine & Growth Factor Reviews 18, 425–433 (2007).

43. Helbig, K. J. & Beard, M. R. The Role of Viperin in the Innate Antiviral Response. Journal of Molecular Biology 426, 1210–1219 (2014).

44. Diamond, M. S. & Farzan, M. The broad-spectrum antiviral functions of IFIT and IFITM proteins. Nat Rev Immunol 13, 46–57 (2013).

45. Deymier, S., Louvat, C., Fiorini, F. & Cimarelli, A. ISG20: an enigmatic antiviral RNase targeting multiple viruses. FEBS Open Bio 12, 1096–1111 (2022).

46. Lv, T., Zhang, J., Zheng, Z. & Fan, X. Single-cell analysis of intermediate horseshoe bat (Rhinolophus affinis) organs reveals viral infections and antiviral immune signatures. BMC Genomics 26, 1011 (2025).

47. Kimura, I. et al. The SARS-CoV-2 Lambda variant exhibits enhanced infectivity and immune resistance. Cell Reports 38, 110218 (2022).

48. Ozono, S. et al. SARS-CoV-2 D614G spike mutation increases entry efficiency with enhanced ACE2-binding affinity. Nat Commun 12, 848 (2021).

49. Kaku, Y. et al. Antiviral efficacy of the SARS-CoV-2 XBB breakthrough infection sera against omicron subvariants including EG.5. The Lancet Infectious Diseases 23, e395–e396 (2023).

50. Kaku, Y. et al. Virological characteristics of the SARS-CoV-2 JN.1 variant. The Lancet Infectious Diseases 24, e82 (2024).

51. Kaku, Y. et al. Virological characteristics of the SARS-CoV-2 XEC variant. The Lancet Infectious Diseases 24, e736 (2024).

52. Motozono, C. et al. SARS-CoV-2 spike L452R variant evades cellular immunity and increases infectivity. Cell Host & Microbe 29, 1124–1136.e11 (2021).

53. Uriu, K. et al. Transmissibility, infectivity, and immune evasion of the SARS-CoV-2 BA.2.86 variant. The Lancet Infectious Diseases 23, e460–e461 (2023).

54. Ozono, S., Zhang, Y., Tobiume, M., Kishigami, S. & Tokunaga, K. Super-rapid quantitation of the production of HIV-1 harboring a luminescent peptide tag. J Biol Chem 295, 13023–13030 (2020).

55. Ozono, S., Zhang, Y., Tobiume, M., Kishigami, S. & Tokunaga, K. Super-rapid quantitation of the production of HIV-1 harboring a luminescent peptide tag. Journal of Biological Chemistry 295, 13023–13030 (2020).

56. Torii, S. et al. Establishment of a reverse genetics system for SARS-CoV-2 using circular polymerase extension reaction. Cell Reports 35, 109014 (2021).

57. Tamura, T. et al. Virological characteristics of the SARS-CoV-2 Omicron XBB.1.5 variant. Nat Commun 15, 1176 (2024).

58. Motozono, C. et al. SARS-CoV-2 spike L452R variant evades cellular immunity and increases infectivity. Cell Host & Microbe 29, 1124–1136.e11 (2021).

59. Reed, L. J. & Muench, H. A SIMPLE METHOD OF ESTIMATING FIFTY PER CENT ENDPOINTS12. Am J Epidemiol 27, 493–497 (1938).

60. Chen, S., Zhou, Y., Chen, Y. & Gu, J. fastp: an ultra-fast all-in-one FASTQ preprocessor. Bioinformatics 34, i884–i890 (2018).

61. Matsuyama, S. et al. Enhanced isolation of SARS-CoV-2 by TMPRSS2-expressing cells. Proc Natl Acad Sci U S A 117, 7001–7003 (2020).

62. Li, H. & Durbin, R. Fast and accurate short read alignment with Burrows-Wheeler transform. Bioinformatics 25, 1754–1760 (2009).

63. Li, H. et al. The Sequence Alignment/Map format and SAMtools. Bioinformatics 25, 2078–2079 (2009).

64. Cingolani, P. et al. A program for annotating and predicting the effects of single nucleotide polymorphisms, SnpEff. Fly 6, 80–92 (2012).

65. Uriu, K. et al. Enhanced transmissibility, infectivity, and immune resistance of the SARS-CoV-2 omicron XBB.1.5 variant. The Lancet Infectious Diseases 23, 280–281 (2023).

66. Dobin, A. et al. STAR: ultrafast universal RNA-seq aligner. Bioinformatics 29, 15–21 (2013).

67. Liao, Y., Smyth, G. K. & Shi, W. featureCounts: an efficient general purpose program for assigning sequence reads to genomic features. Bioinformatics 30, 923–930 (2014).

68. R Core Team. R: a language and environment for statistical computing. Vienna, Austria: R Foundation for Statistical Computing; 2023. https://www.R-project.org/.

69. Love, M. I., Huber, W. & Anders, S. Moderated estimation of fold change and dispersion for RNA-seq data with DESeq2. Genome Biol 15, 550 (2014).

70. Brown, G. R. et al. Gene: a gene-centered information resource at NCBI. Nucleic Acids Res 43, D36–D42 (2015).

71. Wu, T. et al. clusterProfiler 4.0: A universal enrichment tool for interpreting omics data. Innovation 2, (2021).

